# Modulation of the IK_S_ channel by PIP_2_ requires two binding sites per monomer

**DOI:** 10.1101/2021.01.13.426035

**Authors:** Audrey Deyawe Kongmeneck, Marina A. Kasimova, Mounir Tarek

**Affiliations:** Université de Lorraine, CNRS, LPCT, F-54000 Nancy, France

## Abstract

The phosphatidyl-inositol-4,5-bisphosphate (PIP_2_) lipid has been shown to be crucial for the coupling between the voltage sensor and the pore of the potassium voltage-gated K_V_7 channel family, especially the K_V_7.1 channel. The latter, expressed in the myocardium membrane is complexed with its auxiliary subunits, KCNE1 to generate the so-called IK_S_ current. We present here molecular models of transmembrane domain of this complex in its three known states, namely the Resting/Closed (RC), the Intermediate/Closed (IC), and the Activated/Open (AO), robustness of which is assessed by agreement with a range of biophysical data. Molecular Dynamics (MD) simulations of these models embedded in a lipid bilayer including phosphatidyl-inositol-4,5-bisphosphate (PIP_2_) lipids show that in presence of KCNE1, two PIP_2_ lipids are necessary to stabilize each state. The simulations also show that KCNE1 interacts with both PIP_2_ binding sites, forming a tourniquet around the pore and preventing its opening. The present investigation provides therefore key molecular elements that govern the role of PIP_2_ in KCNE1 modulation of IK_S_ channels, possibly a common mechanism by which auxiliary KCNE subunits might modulate a variety of other ion channels.

## Introduction

The IK_S_ current is diffused through the plasma membranes of cardiomyocytes, during the last (fourth) phase of the cardiac action potential (Barhanin et al., 1996; Nerbonne and Kass, 2005; Tristani-Firouzi and Sanguinetti, 1998). This repolarization current is conducted by a tetrameric protein complex derived from the co-expression of four K_V_7.1 α-subunits and KCNE1 ancillary subunits respectively from the KCNQ1 and KCNE1 genes. The K_V_7.1 tetramer forms a voltage-gated potassium (K_V_) channel, a transmembrane protein that upon modification of membrane potential, opens and carries selectively potassium ions to the extracellular medium, while KCNE1 is a transmembrane peptide which acts as an ancillary β-subunit for several K_V_ channels besides the K_V_7.1 one (McCrossan and Abbott, 2004). Together, the α and β subunits are forming the IK_S_ channel complex. K_V_7.1 channels are homo-tetramers of 6 transmembrane helical segments, the first four ones (S1 to S4) forming the voltage-sensor domain (VSD), and the last two (S5 and S6) forming the pore domain (PD) of the channel. As for most K_V_ channels, the VSD and PD are swapped in K_V_7.1. The segment S4 in the VSD carries four positively charged arginines called gating charges that move across the membrane upon depolarization, interacting sequentially with negatively charged side chains from S2 (Wu et al., 2010a). The K_V_7.1 subunit also contains a cytoplasmic C_TERM_ region composed of four cytosolic helices, the first two being connected to the sixth transmembrane segment, the last two being located deeper in the cytosol, forming the tetramerization domain (Wiener et al., 2008). KCNE1 on the other hand is a transmembrane polypeptide of 129 aminoacids divided into one extracellular N_TERM_ domain, a helical transmembrane (TMD) domain, and a cytosolic C_TERM_ domain (Tian et al., 2007).

Numerous mutations of both KCNQ1 and KCNE1 genes are associated with long QT syndromes (LQTS) (Kapplinger et al., 2009; Napolitano et al., 2005; Splawski et al., 2000). The latter are characterized by an extended cardiac action potential that corresponds to the time interval between Q and T waves in electrocardiograms. The LQT phenomenon reflects the inability of the protein complex to generate its IK_S_ outward current and therefore to return cardiomyocytes membranes toward their resting potential. This delay in repolarization disturbs the propagation of the cardiac action potential within the myocardium tissue, and therefore leads to heart rhythm abnormalities, also known as cardiac arrhythmias. Hence, the IK_S_ channel is a therapeutic target for the treatment of LQTS, whose function must be studied and understood to be able to develop any potential effective drug. Over the last twenty years, this protein complex has been extensively studied using various methods (Liin et al., 2015).

The gating of K_V_7.1 and IK_S_ channels is triggered by membrane depolarization and involves three stable states of the VSD: Resting, Intermediate, and Activated. The conformations of these states are known to transition from one to another through the motion of three S4 gating charges, R228 (R1), R231 (R2) and R237 (R4) with respect to two binding sites (Table S1). As in VSDs of most voltage-gated ion channels, that of K_V_7.1 contains two strongly conserved binding sites (residues forming salt bridges with the S4 gating charges): the first one, located in the solvent accessible surface of the VSD, is an acidic residue from S2, E160 (E1). The second one, located deeper in the membrane, is sheltered from the solvent by an aromatic residue from S2, F167, also known as the charge transfer center (CTC), and is composed of two acidic residues, E170 (E2) and D202 (D3) from S2 and S3 segments, respectively (Tao et al., 2010).

These conformational changes also involve a translation of these gating charges through an aromatic residue from S2, F167, which was suggested to constitute the interface between the solvent accessible surface of the VSD, and its occluded site. This residue is conserved in homologous K_V_ channels such as K_V_1.2 or K_V_2.1 (Lacroix and Bezanilla, 2011).

The pore opening is elicited by another mechanism called VSD-PD coupling. The latter occurs in most K_V_ channels and is governed by protein-protein interactions between VSD and PD of distinct α-subunits that couple the activation state of the VSD to the conformation (open or closed) of the PD (Roux, 2006). The exact mechanism for this “electromechanical” process is not completely determined, yet a functional study revealed that the VSD-PD coupling mechanism of the K_V_7.1 tetramer occurs in both the Activated and the Intermediate states of the VSD, but not in its Resting state (Hou et al., 2017). In our recent integrative study, we unveiled the molecular determinants of the distinct intersubunit coupling interfaces that underlie the VSD-PD coupling in the Intermediate and the Activated states of the K_V_7.1 channel (Hou et al., 2020). In presence of KCNE1, this mechanism appears to be hindered when the VSD is in its Intermediate state and enhanced in its Activated state. Hence, the three functional states known for IK_S_ channel are described by the “activation state” of the VSD (R, I or A) and by the conformation (O or C) of the pore, and are referred to as RC, IC and AO states.

The phosphatidyl-inositol-4,5-bisphosphate (PIP_2_) is a membrane phospholipid which participates in the function of many membrane transporters (Hilgemann et al., 2001), including voltage-gated ion channels. PIP_2_ is present at a 1% rate in the inner leaflet of the lipid bilayers forming cell membranes, and experimental studies have shown that it interacts with numerous ion channels including K_V_ channels to modulate their activation. This lipid has been shown to be crucial for the coupling between the VSD and the PD for the K_V_7 channel family (Li et al., 2005; Zaydman and Cui, 2014), especially in the K_V_7.1 channel (Zaydman et al., 2013, 2014).

Hence for instance, the structure of *Xenopus Laevis* K_V_7.1 monomer (KCNQ1_EM_), resolved by cryo-electron microscopy (CryoEM), was locked in a non-physiological conformation of the PD, namely the uncoupled Activated/Closed (AC) state (Sun and MacKinnon, 2017). According to the authors of the study, this might be explained by the absence of PIP_2_, known to be essential for the coupling of the VSD activated state with PD open state in the K_V_7 channels family (Li et al., 2005).

Since beside KCNQ1_EM_ (Sun and MacKinnon, 2017), and the very recently published CryoEM structures of the human K_V_7.1 AO channel in the presence of PIP_2_ (Sun and MacKinnon, 2020), no high-resolution structure of the IK_S_ channel is available, we and others turn to molecular modeling. Quite surprisingly, among the molecular constructs of the IK_S_ channel that have been published the last decade (Gofman et al., 2012; Kang et al., 2008; Ramasubramanian and Rudy, 2018a; Xu et al., 2013), very few were modeled in presence of PIP_2_. Moreover, those which were modeled with the lipid aimed at validating experiments were limited to the identification of the PIP_2_ binding sites (Eckey et al., 2014), without providing any molecular insight about the way the lipid interacts with K_V_7.1 or with its KCNE1 subunits. Recently, a homology model (Jalily Hasani et al., 2017) of the IK_S_ complex was subjected to Molecular Dynamics (MD) simulations in a POPC:PIP_2_ membrane at a 10:1 ratio (Jalily Hasani et al., 2018). Unfortunately, the IK_S_-PIP_2_ interactions were not extensively investigated and the study did not provide information about how PIP_2_ affects the VSD-PD coupling mechanism of the channel. Moreover, despite the fact that several studies have shown that IK_S_ complexes can be expressed in cardiomyocytes with a 4:4 (K_V_7.1:KCNE1) stoichiometry (Murray et al., 2016; Nakajo et al., 2010) most of the IK_S_ models reported so far (Jalily Hasani et al., 2017; Kang et al., 2008; Xu et al., 2013) have not been built with this ratio.

Our previous MD study (Kasimova et al., 2015) of K_V_7.1 models in open and closed states, allowed one to localize the PIP_2_ binding site in the K_V_7.1 subunit, and to characterize the key elements of the K_V_7.1 modulation by PIP_2_ in absence of KCNE1. The lipid was shown to participate in VSD/PD coupling of the K_V_7.1 channel through state-dependent interactions, preventing repulsive forces between basic residues from the S2-S3_LOOP_ and S4 in resting state, and between basic residues from the S2-S3_LOOP_ and S6 in open states. These MD simulations also suggested that PIP_2_ may constitute a third binding site for S4 gating charges. Indeed, the lipid was found to form salt-bridges with R237 (R4) and R243 (R6) in the RC model of K_V_7.1, and not in the AO model. Moreover, the dependence of this lipid for the function of K_V_7.1 was also proved to be increased in presence of KCNE1 due to its additional positive charges in its C_TERM_ domain (Li et al., 2011). This study indicates that IK_S_ may carry an additional PIP_2_ binding site with respect to K_V_7.1. Therefore, the molecular determinants describing a second PIP_2_ binding site in K_V_7.1 subunits in presence of KCNE1 are yet to be investigated. To address these questions, computational chemistry methods are the most insightful ones to unravel the elements of protein function at a molecular level of precision.

Here we propose three models of the K_V_7.1 tetramer in distinct metastable states, in presence of KCNE1 subunits corresponding to the RC, IC and AO states of the IK_S_ channel. Each model has been studied either with one or two PIP_2_ lipids per K_V_7.1 subunit. A total of 6 systems (Figure 1) were submitted each to ~500 ns MD simulations to fully relax the initial constructs. The analyses of these trajectories were then used to assess the validity of the models by comparing them to the literature.

**Figure 1:**
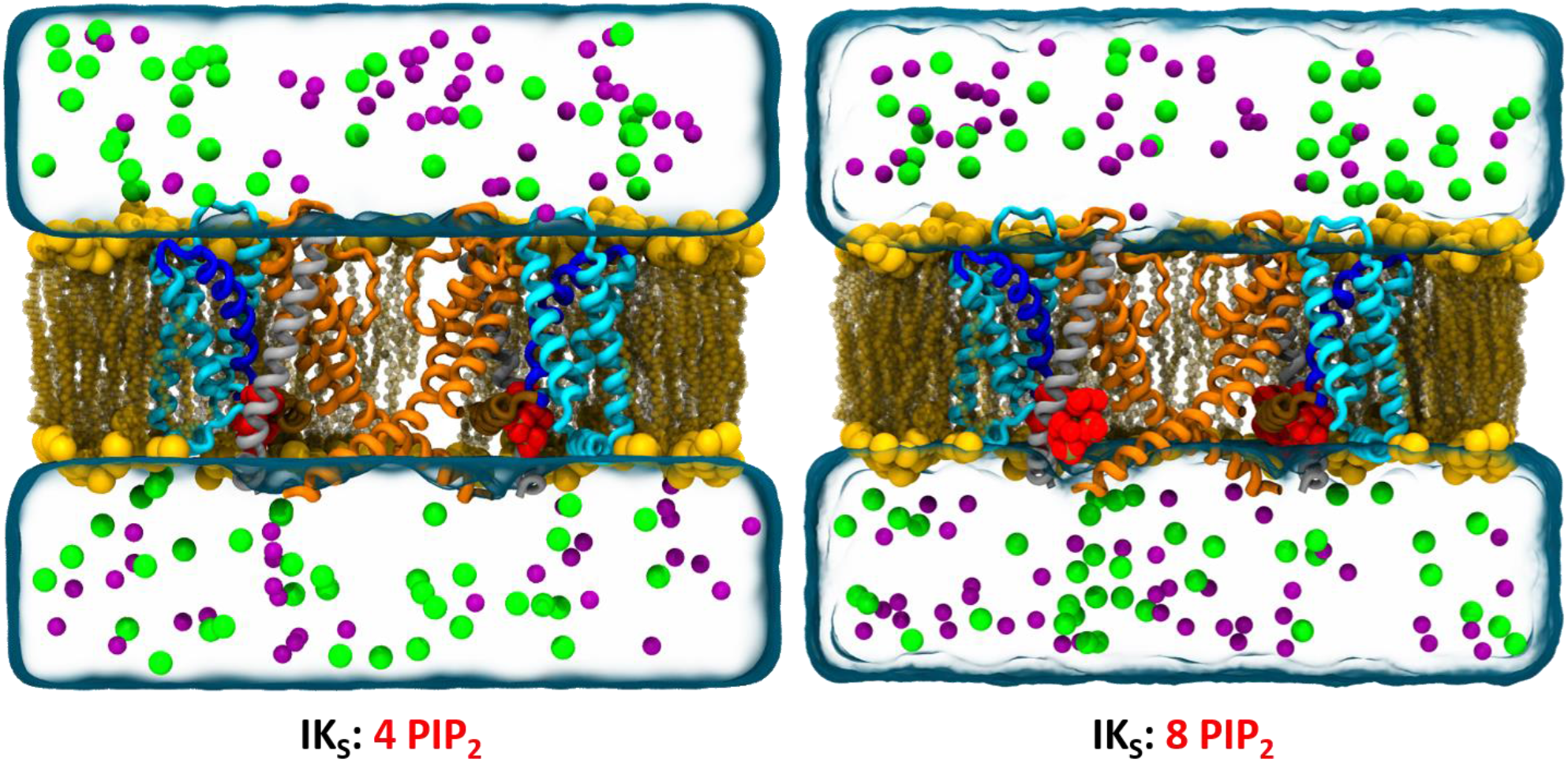
MD systems models of IK_S_ models. Representation of K_V_7.1 models in absence (left) and presence (right) of KCNE1 (gray ribbons) and embedded in a POPC membrane (in yellow spheres) with PIP_2_ lipids (in red spheres). Two K_V_7.1 subunits are shown for clarity. Transmembrane segments are represented in ribbons. VSD segments are colored in cyan, S4 in blue, S4-S5_LINKER_ in brown and PD segments in orange. Membrane-embedded models are surrounded by water (in transparent blue surface) as well as K^+^ (purple spheres) and Cl^-^ (green spheres) ions.

## Results

### Validation of the IK_S_ models: 2 PIP_2_ lipids per subunit are in better agreement with experiments

Molecular modeling methods are based upon the study of representations of chemical and biochemical entities. The main goal of using these computational methods, in the frame of this study, is to gain an atomistic insight on phenomena that were unraveled by experimental studies. Hence, to predict the molecular mechanisms associated to these phenomena, we need to maximize as much as possible the compliance of our models with respect to structural biology data. To achieve this goal, we built our 3D models using structural constraints that were directly drawn from experimental data, and then we monitored the stability of these constraints over the collection of conformations generated by MD simulations, i.e. the trajectories.

To gain a finer insight in the molecular determinants allowing for the stabilization of each state of K_V_7.1 and IK_S_ channels, MD simulations were performed to equilibrate each molecular model within POPC membrane and PIP_2_ lipids. In various K_V_ channels, the activation mechanism of the VSD is mostly characterized by state-dependent salt-bridges between basic residues of S4 and acidic residues spread in S2 and S3 segments (Figure S1). This succession of salt-bridges is describing an upward translation of S4, as well as a clockwise rotation during VSD activation. To validate the state dependent structures of our models VSDs, we monitored the distance between each pair of charge groups that are supposed to interact according to the literature. The state-dependent salt-bridges found in the VSD of our IK_S_ models are reported in Table S1.

In 8 PIP_2_ systems, the interactions between E1 and R228 (R1), and between E2 and R231 (R2), which are specific of the RC state, are both present specifically in our RC model in all subunits. In RC model with 4PIP_2_ system, the interaction between R1 and E1 is present, but the interaction between R2 and E2 is absent, as R2 interacts with E1 in only two subunits out of four. This first result indicates that the presence of two PIP_2_ binding sites in IK_S_ models may be crucial for the stabilization of the VSDs.

In the IC state, S4 translates upwards and rotate clockwise, which leads to E1 interacting with R231 (R2), while E2 interacts with R237 (R4). These interactions have been specifically found in IC model of 8PIP_2_ system, in all subunits for E1-R2 interaction, and in 3 subunits out of 4 for E2-R4 interaction. In the 4PIP_2_ system, E1-R2 interaction is present as expected, but E2-R4 interaction is not absent. Interestingly, R4 also interacts with D3 in all subunits of both systems. The latter forms the charge transfer center along with F167 from S2 and E2. Therefore, R4-D3 interaction may indicate that R4 is fully anchored in the CTC in the 8PIP_2_ system and remain partially anchored in the CTC in the 4PIP_2_ system.

In the AO state, a second upward translation and a clockwise rotation of S4 lead to E1-R4 interaction. This interaction is present only in our AO models of both PIP_2_ systems. In these models, R6 appeared to be anchored in the CTC in all subunits, interacting with both E2 and D3 (data not shown). Noteworthy, the pairs of salt-bridges determined in our IK_S_ models are also present in their respective states of K_V_7.1 models. These MD results highlight the different conformations of the VSD correspond to interaction patterns which remain very stable over time. However, in the 4PIP_2_ systems, state dependent salt-bridge pairs are satisfying experimental data for AO model, but not for IC and RC models.

Overall, these results suggest that IK_S_ models of 8PIP_2_ system are in better agreement with experimental results, as their state dependent expected VSD interactions are more satisfied, and more stable throughout the corresponding MD trajectories. Note that the distinct patterns of gating charge salt-bridges determined in our molecular models support the sliding helix VSD activation mechanism proposed for the K_V_ channels (Durell and Guy, 1992), as well as for the voltage-gated sodium channels (Catterall, 1986).

Besides electrostatic interactions, site-directed mutagenesis can provide important information about the relative position of the KCNE1 helices with respect to the K_V_7.1 subunits. Experiments such as cysteine cross-linking mutagenesis allow one to spot the residues which are close enough to form disulfide bonds. In the framework of such experiments, a disulfide bond can be formed if the Cβ atoms of the corresponding amino acids are within ~ 13 Å (Careaga and Falke, 1992). Thanks to such experimental studies, a significant amount of neighbor residue pairs within the K_V_7.1 channel and the IK_S_ complex have been reported. Accordingly, for each neighbor residue pair, we computed the distance between their respective C_β_ atoms over our MD trajectories and assumed neighbor residue pairs present in the model if the latter was below 13 Å. Among the seven neighbor residue pairs from K_V_7.1 (Table S2), the intersubunit pairs assigned to IC state, as well as the intrasubunit ones assigned to the RC state are present in all IK_S_ models (Figure S2, A), regardless of their respective state and regardless of the amount of PIP_2_ lipids. For the three pairs assigned to the AO state (Figure S2, B), one of those is present in all models, whereas the other two are predominant in most subunits of the AO and RC models, but absent in the IC models. In our K_V_7.1 models, all these pairs are also present in all subunits.

For pairs involving cross linking between K_V_7.1 and KCNE1 residues (Table S3), most of those which have been assigned to the AO state of IK_S_ channel are present in both 4PIP_2_ and 8PIP_2_ AO models (Figure S3). Surprisingly, the only pair of residues assigned to AO state and not present in our AO model (Gln147-Lys41) is present in the RC models only. Among the six residue pairs assigned to the RC state, five of those are present in the 8PIP_2_ system, against four of them in the 4PIP_2_ one. The pairs of residues which are not interacting in RC models are not present in any other model. Finally, for the nine remaining pairs, which were not assigned to any state of the channel, all of them are present in at least one model of each system. Among the three residue pairs that include K_V_7.1 S6 residues and KCNE1 Cterm residues, only one pair is present in both systems. The second pair is present in the 4PIP_2_ system only, and the last pair is absent in both systems.

Overall, K_V_7.1-K_V_7.1 and K_V_7.1-KCNE1 residue pairs are mostly present in all IK_S_ models, regardless of the number of PIP_2_ lipids in the system. Although the presence and the stability of the interactions between each residue pair in our models is testifying for their robustness, these results alone cannot allow to select the best IK_S_ models among their distinct systems.

#### • Neighbor residue pairs: protein-lipid interactions

Besides the protein-protein interactions, several mutagenesis studies conducted on K_V_7.1 in absence and presence of KCNE1 also have highlighted ten basic residues from K_V_7.1 and three others from KCNE1 which can possibly interact with PIP_2_ (Table S4). To characterize these interactions in our IK_S_ models, we calculated average distances between charged groups of each basic residue and PIP_2_. Among the thirteen K_V_7.1 and KCNE1 basic residues which have been shown to be involved in electrostatic interactions with the lipid, ten residues interact with PIP_2_ in at least one MD trajectory of IK_S_ model in 8PIP_2_ system.

Results obtained for these models showed that PIP_2_ inter and PIP_2_ intra are specifically anchored to the VSD and the PD, respectively, in a state independent manner (Figure S4). In each model, residues R190 from the S2-S3_LOOP_ and R249 from the S4-S5_LINKER_ interact with PIP_2_ intra, while residues K362 and R366 from S6 interact with PIP_2_ inter. In addition, some basic residues are also interacting with PIP_2_ in a state dependent manner, such as R192 and R195 from S2-S3_LOOP_, as well as K358 from S6, whose interactions with PIP_2_ are favored in AO model, as well as in IC model for R195. State independent interactions involving VSD residues are related to a motion of PIP_2_, which appears to progressively anchor S2-S3_LOOP_, while remaining bound to S4-S5_LINKER_ residue R249 during VSD activation. State dependent interactions involving PD residues are related to the movement of S6 upon pore opening. PIP_2_ inter progressively binds basic residues of S6 cytoplasmic helices as they spread away from the pore axis towards inner membrane surface where PIP_2_ is localized. Oppositely, the interaction between residue R243 from S4 and the lipid is favored in RC model, which suggests that R243 loses its interaction with PIP_2_ as S4 translates upward during the VSD activation. In IK_S_ models of the 4PIP_2_ system, state-independent interactions between VSD residues and PIP_2_ intra are present, but those between S6 residues and PIP_2_ are merely present. Contrary to K_V_7.1 models in which PIP_2_ intra can bind S6 in AO model, K354 is the only S6 residue that is able to bind PIP_2_ in both IC and AO models of IK_S_ complex. Residues K358, K362 and R366 are located too far away to reach PIP_2_ intra in any of these models.

Four residues are not interacting with PIP_2_ in any IK_S_ system. Residues R181 and K196 from S2-S3_LOOP_ remain too far from PIP_2_ intra throughout MD simulations. R360 from S6, whose sidechains remain tangent to the conduction pathway, was unable to interact with PIP_2_ inter. R259 guanidium group, despite being close to PIP_2_ lipids, cannot get close to the phosphoryl groups of PIP_2_ because of the steric hindrance induced by presence of KCNE1 subunits. Surprisingly, the KCNE1 residues R67, K69 and K70 have their sidechains oriented in two opposite directions, allowing for the ancillary subunit to interact with both PIP_2_ binding sites in a state-dependent fashion in IK_S_ models of 8PIP_2_ system. In RC model, residue K69 binds PIP_2_ intra in two opposite subunits, while R67 and K70 are both binding PIP_2_ inter in all subunits. In IC model, the interaction with K70 is absent, while those with both R67 and K69 are still present. In AO model, K69 is the only KCNE1 residue interacting with PIP_2_. These subsequent interactions over all our stable state models of IK_S_ of 8PIP_2_ system suggest that KCNE1 undergoes two clockwise rotations during VSD activation, each occurring during the RC-IC and the IC-AO transitions, respectively. In the IC and RC models, KCNE1 residues are binding both inter and intra PIP_2_, forming a circle around the cytoplasmic region of S6 helices. This PIP_2_-KCNE1 circle was not observed in the AO model, as KCNE1 does not bind PIP_2_ intra. Thus, the cytoplasmic region of S6 can spread away from the pore axis towards PIP_2_ inter. Quite interestingly, in the MD trajectories of IK_S_ models in the 4PIP_2_ system, a likewise PIP_2_-KCNE1 circle cannot be formed due to the absence of PIP_2_ inter. Indeed, interactions involving R67 and K70 are impaired, while those involving K69 are conserved. However, in the AO model, the interaction between K69 and PIP_2_ is impaired, and basic residues of KCNE1 are facing those of S6 that strongly interact with PIP_2_ inter in 8PIP_2_ systems, which might generate electrostatic repulsions, leading to the collapse of S6 segments and pore closure.

In summary, our results highlighted an additional PIP_2_ binding site in IK_S_ channel, with which KCNE1 interacts in a state dependent manner. This additional PIP_2_ binding site allow for the shaping of a PIP_2_-KCNE1 circle in the uncoupled RC and IC states model of IK_S_ channel. This may allow for the control of S6 helices conformations by the KCNE1 subunits.

To verify this hypothesis, results of pore radii calculations in our IK_S_ models turn out to be useful. Indeed, results obtained for the IK_S_ models will allow to figure out if the presence of PIP_2_ inter in the 8PIP_2_ systems is inducing a change in the width of the conduction pathway.

#### • Pore radii

Pore radii calculations have been conducted to select the best IK_S_ model among the systems with 4 PIP_2_ lipids and the ones with 8PIP_2_ lipids. Based on the correlation identified between the pore size of a K_V_ channel and the free energy associated to the conduction of a potassium ion (Treptow and Tarek, 2006), the analysis of the pore size of our models can allow to assess the reliability of our IK_S_ models with respect to those of K_V_7.1 model, and also with respect to experimental data. Integrative studies conducted on the Shaker channel (Díaz-Franulic et al., 2015) highlighted the fact that the cytoplasmic region of K_V_ channel conduction pathway need to be able to accommodate a solvated potassium ion, whose minimal radius is ~3.6 Å, to be open. Pore radius measurement of the conduction pathway of KCNQ1_EM_ structure (Sun and MacKinnon, 2017) in its Activated/Closed state reveals three S6 regions of constriction in the cytoplasmic side of the conduction pathway: the upper constriction is formed by the backbones of G345; the middle constriction is formed by hydrogen bonds between S349 hydroxyl groups, and the lower constriction is formed by the L353 side chains that are oriented towards the center of the pore and binding each other to form a hydrophobic seal. These three regions of constriction present pore radii of ~2 Å for G345, ~0.8 Å for S349, and ~1.15Å for L353, which may prevent a K^+^ ion to go through this pathway, as its ionic radius is ~1.33Å. This pore radius profile provides a hint on the constriction zones of K_V_7.1 pore when the channel is decoupled due to the absence of PIP_2_ lipids. Hence, we compared first the pore radii profiles of IK_S_ MD trajectories in its distinct systems, and then we compared the most robust model of IK_S_ in its three states with those of K_V_7.1 models.

We mapped the average pore radii to see if these residues are forming a constriction in our IK_S_ models. The MD trajectories of our models suggest that the backbone of G345 is oriented toward the center of the pore, while the side chains of both S349 and L353 remain tangent to the pore surface in all models. Nevertheless, the average pore radii estimated on these regions (Figure 2) shed light on state dependent opening / closure of the conduction pathways. The models indicate, in the AO models (Figure 2A), values are between 2.7 Å and 5 Å. In the Intermediate models (Figure 2B), these values are between 2 Å and 4.5 Å. In the resting models (Figure 2C), the average pore radii at the level of constricted regions assume values between 1 Å and 3 Å for the three models.

**Figure 2:**
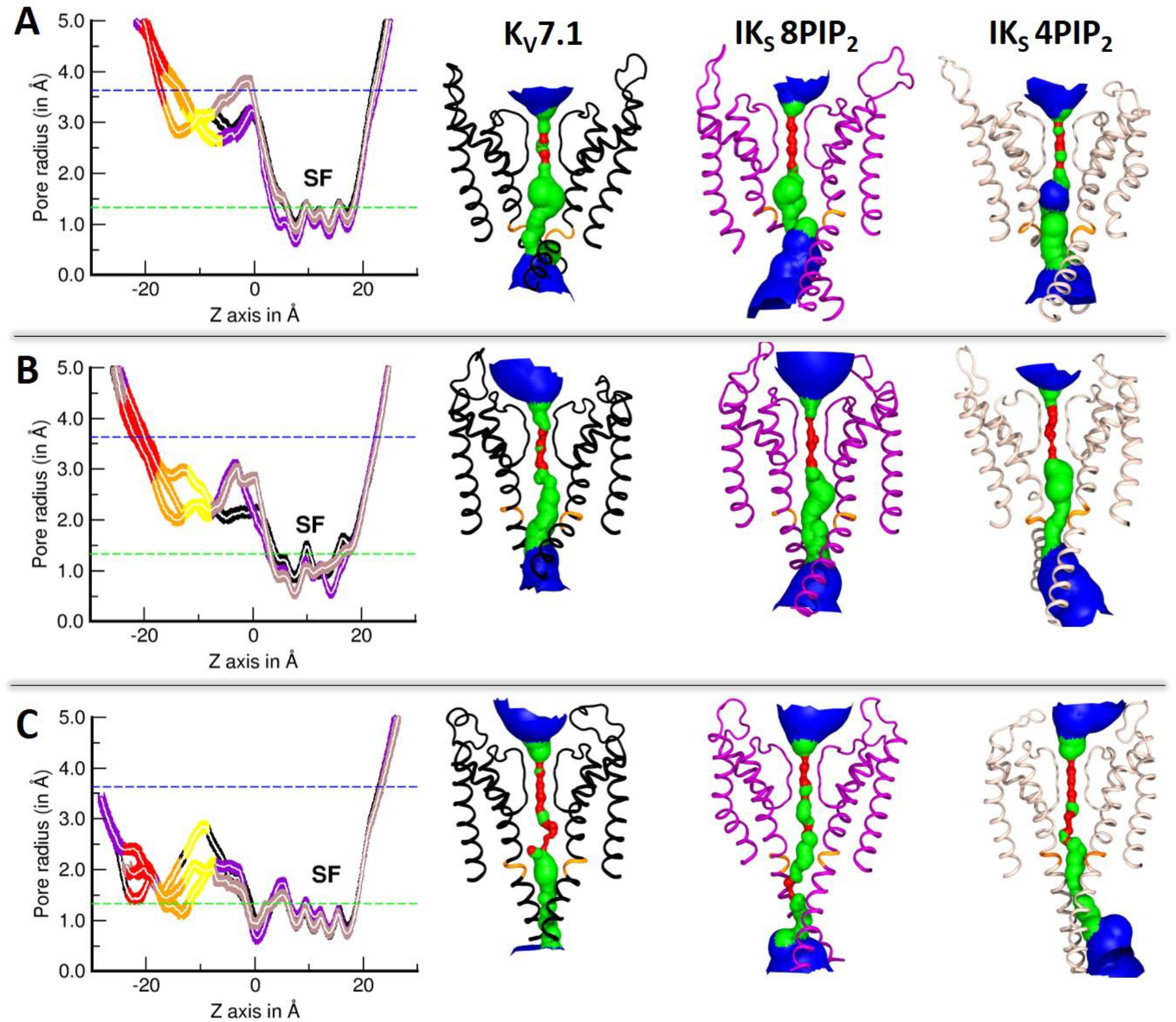
Validation of IK_S_ models through pore radii calculations. The graphs report the average pore radii (left) along the conduction pathway of K_V_7.1 (black curves) IK_S_ 8PIP_2_ (purple curves) and IK_S_ 4PIP_2_ (beige curves) models in **A.** AO state **B.** Intermediate state and **C.** RC state. Averages radii at the levels of residues G345, S349 and L353 are depicted in yellow, orange and red curves, respectively. The right panels show cartoon representations of K_V_7.1 (black) IK_S_ 8PIP_2_ (purple) and IK_S_ 4PIP_2_ (beige) models. Potassium ionic (1.33 Å) and hydrodynamic radii (3.6 Å) are represented by green and blue dashed lines in the graphs, respectively. Pore solvent accessible surfaces are colored as follows. Pore radii values inferior to K+ ionic radius, are colored in red. Pore radii values ranging between K+ ionic radius and K+ hydrodynamic radius, are colored in green. Pore radii values superior to K+ hydrodynamic radius, are colored in blue.

To discriminate one IK_S_ model from another, we compared for each state, the average pore radii values obtained at the level of the constriction zones formed by G345, S349 and L353 in IK_S_ models with those of KCNQ1_EM_ structure. In the AO state, K_V_7.1 ionic current is increased in presence of KCNE1, thus IK_S_ AO models should present wider pore radii than KCNQ1_EM_ structure at the level of constriction regions. Among our AO models (Figure 2A), the IK_S_ 8PIP_2_ system have the least constrictions. Pore radii values of ~ 3 Å, 3.9 Å and 4.5Å in average at the levels of G345, S349 and L353, were respectively found. The IK_S_ 4PIP_2_ AO model in contrast appears to be more constricted at the level of S349, as its average pore radii is 0.9 Å lower than those of other AO models. A decrease of 1 Å in the radius value of a nanopore can lead to a significant rise in the free energy cost necessary for conduction (Díaz-Franulic et al., 2015; Treptow and Tarek, 2006). This result, in addition to the number of protein-lipid interactions we found in IK_S_ AO models, suggest that the IK_S_ AO model is in better agreement with experimental data when embedded in 8PIP_2_ system than in 4PIP_2_ system.

In the Intermediate state, the ionic current is abolished in presence of KCNE1, so IK_S_ models should present a smaller pore radius than in K_V_7.1. In the intermediate models (Figure 2B), the pore radii of the constriction zone of L353 are ~3.6 Å in all models. Therefore, we only considered G345 and S349 constriction zones. The K_V_7.1 model present the highest average pore radii values, (~ 2.7 Å and 2.9 Å at the levels of G345 and S349, respectively) compared to the IK_S_ models (both ~ 2 Å). Hence, in a nutshell, the results obtained for intermediate models suggest that K_V_7.1 conduction pathway is ~0.8 Å narrower in presence of KCNE1, regardless of the number of PIP_2_ lipids in the system which agrees with experiments.

In summary, constriction zones of RC models (Figure 2C), present pore radii values below the minimal hydrodynamic radius of K^+^ ions of 3.6 Å, which suggest that in RC models, the pore is closed regardless of the presence of KCNE1 or the number of PIP_2_ lipids.

The differences observed between the average pore radii calculated over the MD trajectories obtained for K_V_7.1 and IK_S_ models indicate that KCNE1 may increase the tightening of the inner pore in the RC and IC states, while inducing a closer proximity between S6c region and the inner membrane surface in the AO state, which agrees with the experimental studies. The comparison of the average pore radii obtained for the IK_S_ models of both 4PIP_2_ and 8PIP_2_ systems with those obtained for the K_V_7.1 models indicate that the MD trajectories of the IK_S_ models of 8PIP_2_ system fit better the ionic conductance measures of the IK_S_ channel (Wu et al., 2010b) compared to the K_V_7.1 channel.

## Discussion and Conclusion

The present work provides tridimensional models of three states of the K_V_7.1 channel in presence of KCNE1, whose features are mostly satisfying the structural constraints drawn from experimental studies. Our previous computational study of K_V_7.1 models (Kasimova et al., 2015) highlighted two components of VSD-PD coupling: a protein-protein component and a protein-lipid component. Protein-protein components of this mechanism were mostly characterized by electrostatic interactions between residues from the S4-S5_LINKER_ and residues from S6, while the protein-lipid components were characterized by state-dependent interactions between PIP_2_ and the K_V_7.1 subunits.

In our IK_S_ models, considering a second PIP_2_ binding site per subunit was required in order to optimize the agreement with experimental results. This work shows that in contrast to the case of K_V_7.1, in IK_S_ models, PIP_2_ lipids engage in state-independent interactions with K_V_7.1 subunits: PIP_2_ intra, which is present in both K_V_7.1 and IK_S_ models, predominantly binds the lower VSD basic residues, while PIP_2_ inter, presumably most required in IK_S_ channels, binds the basic residues from S6 and KCNE1. In each subunit, PIP_2_ inter is located between C_TERM_ regions of both the KCNE1 helix and the S6 segment (Figure S4). In 8PIP_2_ IK_S_ models (Figure S4, A, upper panel), PIP_2_ inter remain bound to the S6 residues K362 and R366 in a state independent manner, as well as to the KCNE1 residues R67 K69 and K70 in a state dependent manner. In the 4PIP_2_ IK_S_ AO system, which is lacking the PIP_2_ inter (Figure S4, B, upper panel) the basic residues from S6 and from KCNE1 might repel each other and prevent S6 Cter from reaching PIP_2_ intra located in the inner membrane surface in AO state, as observed.

The molecular determinants of this second PIP_2_ binding site we identified in our IK_S_ models are supported by a previous functional study (Li et al., 2011) which shed light on several residues of KCNE1 subunit participating in KCNE1-PIP_2_ interactions in IK_S_ channels. Our results are also in agreement with an integrative study including both experimental and computational approaches (Eckey et al., 2014), which aimed at identifying K_V_7.1 interactions with PIP_2_ in IK_S_ that highlighted the existence of two PIP_2_ binding sites for this complex, one for VSD residues and a second one for PD residues.

Noteworthy, KCNE1 residues bind both PIP_2_ inter and PIP_2_ intra in the RC and IC states which may induce a tighter packing of PD segments. Indeed, these KCNE1-PIP_2_ interactions form a circle of electrostatic interactions around the cytoplasmic region of S6 (Figure 3A, B). This circle may act as a tourniquet (Figure 3A, B, top panels), inducing a shrinkage of the conduction pathway, leading to the lower pore radii values we obtained for IK_S_ RC and IC models of 8PIP_2_ with respect to IK_S_ RC and IC models of 4PIP_2_, respectively. In the AO models of IK_S_, KCNE1 loses its interactions with PIP_2_ intra and binds only PIP_2_ inter (Figure 3C). Through this interaction, one can predict that KCNE1 is pulling S6 helices towards the inner leaflet of the membrane, as PIP_2_ inter remains bound to S6 basic residues.

**Figure 3:**
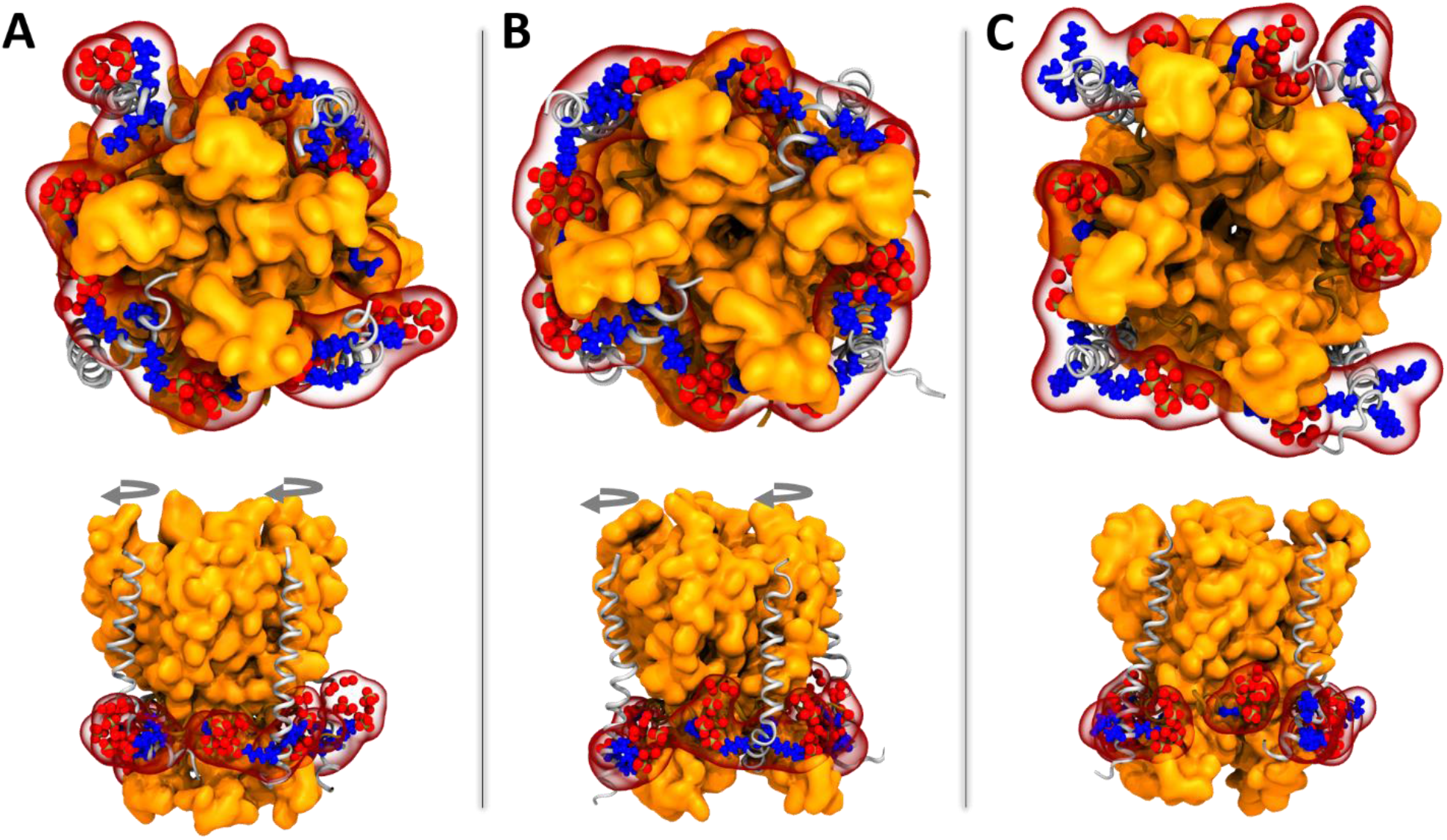
Structural mapping of KCNE1 basic residues and PIP_2_ lipids in the pore domain of IK_S_ models in 8PIP_2_ systems. Intracellular view (top panels) and side view (bottom panels) of K_V_7.1 pore domain segments (in orange surface) along with S4-S5_LINKER_ (in brown ribbons), KCNE1 subunits (in gray ribbons) and PIP_2_ phosphate groups (in red spheres) in **A.** RC model, **B.** IO model and **C.** AO model. KCNE1 basic residues R67, K69 and K70 are shown in blue spheres. S4-S5_LINKER_ residue R249 that interact with PIP_2_ intra in a state independent fashion is shown in blue sticks. The successive rotational movements of KCNE1 subunits predicted to occur during RC-IC and IC-AO transitions are shown in gray circled arrows.

In the RC and IC models, the KCNE1 basic residues form electrostatic interactions with both PIP_2_ binding sites. This network of interactions forms a circle around S6 Cterm helices which seems to prevent those helices from moving away from the pore axis and therefore preventing the conduction pathway of the IK_S_ channel from expending. Nevertheless, these pore radii differences between the studied IK_S_ models estimated from the MD trajectories did not allow us to determine if these models are genuinely able to conduct potassium ions. Further studies allowing for the calculation of the free energies associated to ion translocation, beyond the scope of this work, would be required to address this issue.

As a matter of fact, a recent computational study, which consisted in the use of machine learning methods to generate a conformational space of IK_S_ channels, have yielded to the design of a structure-based predictor of IK_S_ channel experimental properties including its subconductance and gating current (Ramasubramanian and Rudy, 2018b). The two sequential translations and rotations of S4 and the rotations of KCNE1 leading to VSD activation we predicted from our models are supported by the results obtained with this structure-based predictor.

The results reported by Li et al. (Li et al., 2011) have shown that KCNE1 increases PIP_2_ sensitivity 100-fold over channels formed by the pore-forming K_V_7.1 α-subunits alone. In this study the authors identified four residues (R67, K69, K70, and H73) in proximal C-terminus of KCNE1 as key determinants of PIP_2_ sensitivity. Mutations of these key residues in KCNE1 (R67C, R67H, K70M, and K70N) are associated with long QT syndrome (Hedley et al., 2009; Kapplinger et al., 2009; Lai et al., 2005). They reduce IK_S_ currents and PIP_2_ sensitivity. Application of exogenous PIP_2_ to these mutants restores wild-type channel activity. The results reported in the study of Li et al reveal the vital role of PIP_2_ for KCNE1 modulation of IK_S_ channels, confirming the previous studies that highlighted the inhibitory effects of PIP_2_ membrane depletion on IK_S_ channel function, by inducing PIP_2_ depletion through the cotransfection of a IK_S_ channel construct (Dahimène et al., 2006; Wang et al., 1998) and a PIP_2_-phosphatase in Human Embryonic Kidney (HEK-293) cells (Royal et al., 2017; Suh et al., 2006). Furthermore, other studies reported that PIP_2_ acts as a second messenger (Logothetis et al., 2010) of various ion channels. In the case of IK_S_ channel, PIP_2_ participates in the transduction of sympathetic signalling pathways induced by stress (Matavel and Lopes, 2009) or exercise (Dvir et al., 2014), leading to a left-shift on IK_S_ voltage-dependence of activation (O-Uchi et al., 2015) and to a 2-fold increase of IK_S_ current amplitude (Marx et al., 2002), respectively.

The increase in PIP_2_ membrane levels appeared to restore the function of several loss-of-function mutations (R174C, R243C and R336Q) of KCNQ1 gene (Park et al., 2005) that are related to impaired sympathetic stimulation pathways of IK_S_ channel (Matavel et al., 2010). Specifically, this observation suggests that both sympathetic pathways mutants activate the IK_S_ channel by strengthening its interactions with VSD residues R174 and R243 as well as with PD residue R366. The present investigation, confirming that two of these residues are involved in specific interactions with PIP_2_, provides the key molecular elements that govern such a role. Indeed, our models suggest that residues R243 and R366 bind PIP_2_ intra and PIP_2_ inter in a state-dependent and a state-independent fashion, respectively. This observation agrees with the sympathetic stimulations of the VSD mutants and PD mutants cited above which yielded to different sensitivities to PIP_2_ membrane levels. Altogether, these results indicate a possible difference of binding affinity between both PIP_2_ binding sites in IK_S_ channel.

Moreover, the family of ion channel β-subunits (KCNE1-5) contains several members that have been reported to modulate the activity of a variety of channel α-subunits in ion channel complexes. Many of these channel α-subunits or channel complexes are also modulated by PIP_2_. The KCNE1 basic residues listed above that are essential for modulation of the IK_S_ PIP_2_ sensitivity are highly conserved across all members of the KCNE family of peptides (Figure 4), suggesting that modulation of PIP_2_ sensitivity may be a common mechanism of current modulation by the KCNE β-subunits. As we built models of the transmembrane regions of both K_V_7.1 and KCNE1 subunits, the recently reported PIP_2_ binding sites, located in the cytoplasmic region of the K_V_7.1 (Tobelaim et al., 2017) and the K_V_7.3 subunits (Choveau et al., 2018), could not have been addressed in the frame of this study.

**Figure 4:**
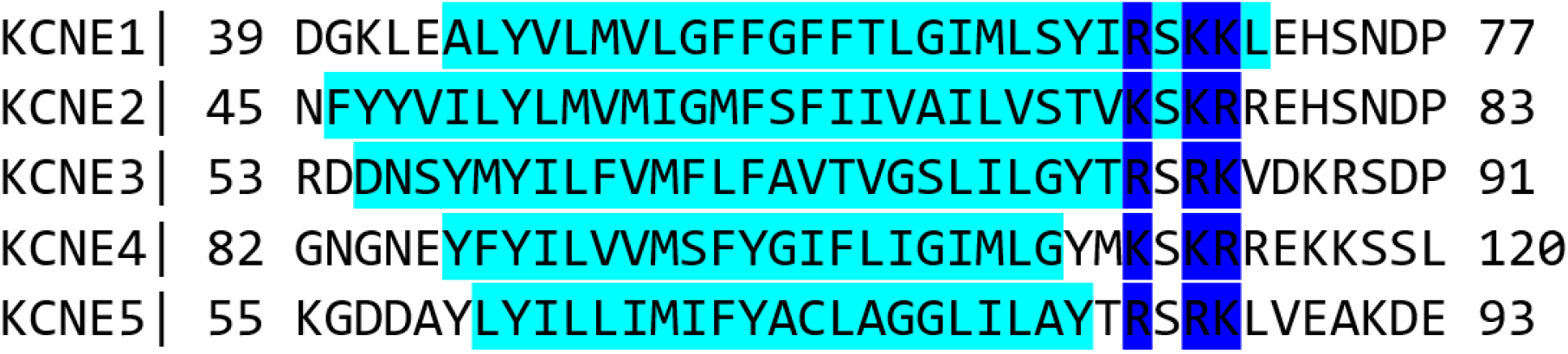
Conserved basic residues in the of KCNE ancillary subunits. The picture shows a sequence alignment of the transmembrane domains (TMD) of KCNE subunits, highlighted in cyan. The conserved basic residues, located at the end of the TMD, near their C_TERM_ domains, are highlighted in blue.

The present analysis demonstrated the coherence of the simulation of our IK_S_ models in a system that contain eight PIP_2_ molecules instead of four, each located in the respective binding sites of K_V_7.1 and KCNE1 subunits, in order to fit the experimental data as much as possible. Hence, the robustness of the resulting MD trajectories will be used for ongoing work that features the investigation of the molecular determinants of the VSD-PD coupling and pore opening mechanisms of the K_V_7.1 channel, in both the absence (Hou et al., 2020) and the presence of the KCNE1 ancillary subunit.

## Material and Methods

To investigate the structural determinants of PIP_2_ binding sites of the IK_S_ channel, we first needed to build molecular models which had to be as trustworthy as possible with respect to experimental data available. Since no high-resolution structure of the K_V_7.1 channel was available yet, we used homology modeling to build several models of the channel embedded within a lipid bilayer (Page et al., 2007). For K_V_7.1, the activation mechanism involves three stable states, i.e., RC, IC, and AO states. Accordingly, each state was modeled to obtain a larger spectrum of possible α-subunits conformations, allowing for the prediction of possible transition mechanisms for IK_S_ channel. For these models, the N_TERM_ and C_TERM_ cytoplasmic regions of K_V_7.1 were ignored. Only residues 122 to 366 from KCNQ1 human sequence, corresponding to the transmembrane region of the channel, were considered. To fully characterize the modulation of K_V_7.1 by the KCNE1 subunits, we built each state model along with the TMD (residues 39 to 76) of the human KCNE1 NMR structure (Tian et al., 2007) using a K_V_7.1:KCNE1 subunit ratio of 4:4.

### • Homology modeling of K_V_7.1 in its distinct states

Homology modeling, also known as comparative modeling, aims at building a protein structure from its primary sequence, starting from the premise that two proteins with similar primary sequences will be displaying similar folds (3D structures). To adjust the salt-bridge patterns of the VSD with respect to experimental results (Wu et al., 2010b, 2010a), and thereby obtain distinct activation state models of the VSD, the charged group of E160 (E1) was constrained to be in close proximity with:

– R237 (R4) in activated/open (AO) model, using the refined crystallographic structure of K_V_1.2 (Chen et al., 2010) as a template
– R231 (R2) in Intermediate/Closed (IC) model using the γ conformation of the K_V_1.2 subunit refined structure obtained from previous MD simulations (Delemotte et al., 2011, 2015)
– R228 (R1) in Resting/Closed (RC) model, using the ε conformation of K_V_1.2 refined structure obtained from the aforementioned study.

The alignment of the K_V_7.1 human sequence and K_V_1.2 rat sequence was first conducted automatically, using ClustalW2 (Larkin et al., 2007). For the PD, the percentage of sequence identity between K_V_7.1 and K_V_1.2 is 36%. For the VSD, the percentage is 19.5%. This lower sequence identity is mainly due to the S2-S3 loop, which is longer in K_V_7.1 sequence than in the one of K_V_1.2. To overcome this discrepancy, this alignment was refined manually and locally. Indeed, we specifically aligned K_V_7.1 important residues (conserved acidic residue of S2, conserved CTC, S4 conserved gating charges) with similar K_V_1.2 ones. Plus, insertions and deletions were concentrated in the loop regions. Eventually, without S2-S3 loop, the percentage of identity between K_V_7.1 and K_V_1.2 was increased to 25% for the VSD.

For each stable state of K_V_7.1, fifty models were generated using MODELLER (Eswar et al., 2007). This software performs constrained modeling, a technique which consists in using template coordinates and sequence alignment information as constraints for the building of 3D models. S2-S3 loop has been modeled from a template 3D structure extracted from NMR data (Peng et al., 2014), which suggests a helical structure for this connecting loop. Since MODELLER allows one to add specific geometric restraints, several ones were applied according to site-mutagenesis results to increase the reliability of these models with respect to experimental data. Other constraints drawn from 3D models of IK_S_ channel stable states (Kang et al., 2008) were also used to predict the position of KCNE1 subunits with respect to K_V_7.1.

Among these 50 obtained models, the best 10 were selected according to their potential energy values, calculated using DOPE (Discrete Optimizing Protein Energy) (Shen and Sali, 2006) knowledge-based scoring function, implemented in MODELLER. The stereochemical quality of these models were evaluated using PROCHECK software (Laskowski et al., 1993). For each stable state, the structure presenting the highest number of Phi and Psi torsion angles in Ramachandran’s plot well favored areas (>95%), as well as the lowest number of torsion angles in the disfavored areas (<5%), was chosen to perform molecular dynamics simulations to study our models.

### • Molecular Dynamics simulations

To reproduce the behavior of IK_S_ channels in their natural environment, the three models were embedded in lipid bilayers prior to the simulations. To do this, we used a method available in the input generator of CHARMM (Jo et al., 2008), which consists in adding lipid molecules around the protein structure (Wu et al., 2014). Since Phosphatidylcholine (PC) lipids are the most abundant lipids found in cell membranes, Palmitoyl-Oleyl PC (POPC) lipids were selected to build the bilayer in each of our nine systems. To incorporate phospholipids in the correct binding sites of the channels, a PIP_2_ molecule was added within the inner leaflet of the bilayer, at the bottom of each VSD (PIP_2_ intra) with respect to experimental studies (Eckey et al., 2014; Zaydman et al., 2013) and computational results of MD simulations conducted on K_V_7.1 subunits along with PIP_2_ (Kasimova et al., 2015; Zaydman et al., 2014). We embedded the RC, IC and AO states models in two distinct environments. In the first system, referred to as 4PIP_2_, we added only PIP_2_ intra, corresponding to the binding site of K_V_7.1 subunits. In the second system, referred to as 8PIP_2_ we added a second PIP_2_ molecule at the bottom of KCNE1 subunits, following data from experiments that highlighted a second PIP_2_ binding site in IK_S_ channels (Li et al., 2011) (cf. Figure 1.)

Simulations were carried out using the NPT ensemble for the equilibration of the systems, at 300K, and 1 atm. (Noteworthy, POPC has a transition temperature above 300K, *i.e.* it is in its liquid crystal phase). Chemical bond lengths between hydrogens and heavy atoms were all constrained at their equilibrium values so that a time-step of 2.0 fs could be used. The systems (lipids+channel) were surrounded by a 150 mM [KCl] solution. We used the CHARMM36 force-field (Huang and Mackerell, 2013), along with CMAP correction (Buck et al., 2006) and NAMD code (Phillips et al., 2005) to perform all MD calculations.

The MD simulations were conducted in four steps, during which motion constraints were applied on the whole system, and then gradually released. The first step of 200 ps aimed at fully solvating the protein in the membrane, by letting water molecules rearrange themselves around the protein. Accordingly, constraints were set up on all IK_S_ atoms. The second step of 6ns was run to relax the side chains of the protein, so the constrains were kept only on IK_S_ backbone. During these two steps, the positions of PIP_2_ phosphorus atoms of in the system were kept constant, to maintain this lipid in its correct binding sites. The third step was conducted to allow PIP_2_ lipids to rearrange around the protein complex and within the lipid bilayer. Hence the constraints on PIP_2_ phosphorus atoms were removed but we maintained on the backbone of IK_S_ complex. This step of 70 ns was also conducted to let the density of the system reach a constant value. Finally, the last step corresponds to the so-called production phase. This step, performed without any specific forces on any coordinate of the system, lasts approximately 500 ns. Only the backbone of the selectivity filter (corresponding to the voltage-gated ion channel conserved 311-TTIGYG-316 sequence) was spatially constrained, to prevent ion conduction during the simulations.

### • Validation of IK_S_ models by MD simulation analyses: Strategy

We considered here only the production phase of MD simulations. To assess the reliability of the models with respect to experimental data, we confronted, for each state of the IK_S_ channel, the 2 systems against a set of results obtained from biophysical characterization. To achieve this, we first gathered all the cross-linking studies conducted on IK_S_ channel to identify: (i) The VSD salt-bridge patterns, (ii) The K_V_7.1 intersubunit and intrasubunit neighbor residues, (iii) The KCNE1/K_V_7.1 neighbor residues and (iv) The K_V_7.1 and KCNE1 residues which bind PIP_2_. Cysteine cross-linking studies assume that mutated cysteine residues can form disulfide bonds with a significant formation rate constant if the distance between their Cβ atoms is ≤ 13.2 Å (Careaga and Falke, 1992). Accordingly, we evaluated the Cβ-Cβ distances of each pair of residues in the MD trajectories we obtained for our IK_S_ models and reported their average values (see tables S2-S3 (Supporting Information)).

A pair was considered as fulfilled in the model if the average distance between the Cβ atoms pair was below ~ 13 Å for at least three channel subunits out of four. These distances were monitored every 2 ns over the MD trajectories, using a designed program written in TCL language which we executed within the scripting interface of Visual Molecular Dynamics (VMD) software (Humphrey et al., 1996). We investigated the electrostatic interactions and the hydrogen bonds that are expected between the S4 gating charges and the S2 negative ones (Table S1), as well as the ones between basic side chains from the cytoplasmic region of KCNE1 and PIP_2_ (Table S4). These were monitored every 2 ns and were considered satisfied if the average distance between the respective charge moieties were below 3.5 Å, which is the average distance encountered for ion pairs in protein NMR structures (Kumar and Nussinov, 2002).

For these calculations, we considered that the positive charge of arginine side chains is delocalized between three terminal nitrogen atoms of its guanidium group, and the negative charges of glutamate and aspartate side chains are shared by two oxygen atoms from their respective carboxyl group. For PIP_2_, its five negative charges are delocalized between eight oxygen atoms from its three phosphate groups (Figure S2). Hence, the salt bridges they form with the channel residues were monitored by computing each of the atom pair combinations of terminal nitrogen atoms from Arg guanidium group and terminal oxygens from Glu or Asp carboxyl groups. The distance graphics were designed using R scripting language (Heiner Schwarte et al., 2017). For each pair of residues, an interaction was considered present if found in at least three subunits out of four.

Starting from the premise that pore radius of open K_V_ channels gates energetically favors ionic conduction (Beckstein et al., 2004; Peter and Hummer, 2005; Treptow and Tarek, 2006), one expects that channel models with larger pore radii are likely to facilitate ion conduction and therefore generate similar ionic currents as those recorded experimentally. Pore radii and pore solvent accessible surfaces of all models were calculated and generated every 20 ns of their respective MD trajectory, both using HOLE program (Smart et al., 1996). For each model, we reported the average pore radii values along the pore axis, i.e. the conduction pathway. For each state, we compared the obtained pore radii with those calculated for the K_V_7.1 models built in the frame of a previous study. Indeed, KCNE1 is known to enhance K_V_7.1 ionic currents in the AO state, while abolishing this current in both the Intermediate and the RC states. Pore surfaces were rendered with VMD.

For each IK_S_ state, the MD trajectory which presented the highest number of salt bridges in agreement with experimental data, and the most relevant average pore radii with respect to those obtained for the K_V_7.1 models, was selected as the best model of IK_S_ channel complex.

## Supplementary Material

**Table S1:** State dependent VSD salt bridges in IK_S_ models. Identification of K_V_7.1 state dependent salt-bridges of S4 gating charges, determined through charge reversal mutagenesis, in our IK_S_ models. Each residue name is colored according to the chemical nature of its sidechain. Basic residues in blue, acidic residues in red, and polar residues in green.

| Charge reversal mutants | References | State | Number of subunits in 4PIP<sub>2</sub> systems (over 4) |  |  | Number of subunits in 8PIP<sub>2</sub> systems (over 4) |  |  |
| --- | --- | --- | --- | --- | --- | --- | --- | --- |
|  |  |  | AO | IC | RC | AO | IC | RC |
| **E160R/R237E** | (Wu et al., 2010a; Zaydman et al., 2014; Wu et al., 2010b) | AO | 4 | 0 | 0 | 3 | 0 | 0 |
| **E160R/R231E** |  | IC | 0 | 4 | 2 | 0 | 4 | 0 |
| **E170R/R237Q** |  | IC | 0 | 2 | 1 | 0 | 3 | 0 |
| **E160R/R228E** |  | RC | 0 | 0 | 4 | 0 | 0 | 4 |
| **E170R/R231E** |  | RC | 0 | 0 | 1 | 0 | 0 | 4 |

**Table S2:** State-dependent protein-protein interactions involving K_V_7.1 residues. Identification of the K_V_7.1 neighbor residue pairs previously identified through site-directed mutagenesis in our IK_S_ models. The residues involved in intrasubunit interactions are marked with an asterisk. The remaining residue pairs are engaged in intersubunit interactions. Each residue name is colored according to the chemical nature of its sidechain. Apolar residues (aliphatic, aromatic) are written in black, and polar residues in green.

| K<sub>v</sub>7.1 residue pairs |  |  |  | State | Reference | Number of subunits in 4PIP<sub>2</sub> systems (over 4) |  |  | Number of subunits in 8PIP<sub>2</sub> systems (over 4) |  |  |
| --- | --- | --- | --- | --- | --- | --- | --- | --- | --- | --- | --- |
| Residue 1 |  | Residue 2 |  |  |  | AO | IC | RC | AO | IC | RC |
| **Thr144** | S1 | **Tyr299** | P-helix | AO | (Xu et al., 2013) | 4 | 4 | 4 | 4 | 4 | 4 |
|  |  | **Ser298** |  |  |  | 4 | 1 | 4 | 4 | 0 | 4 |
|  |  | **Ala300** |  |  |  | 4 | 0 | 4 | 4 | 0 | 3 |
| **Phe232** | S4 | **Phe279** | S5 | IC | (Nakajo and Kubo, 2014) | 4 | 4 | 3 | 4 | 4 | 3 |
| **Leu250\*** | S4-S5<sub>L</sub> | **Leu353\*** | S6 | RC | (Choveau et al., 2011) | 4 | 4 | 4 | 4 | 4 | 4 |
| **Val254\*** |  |  |  |  |  | 4 | 4 | 4 | 4 | 4 | 4 |
| **His258\*** |  |  |  |  |  | 4 | 4 | 4 | 4 | 4 | 3 |

**Table S3:** State-dependent protein-protein interactions involving K_V_7.1 and KCNE1 residues. Identification of the K_V_7.1-KCNE1 neighbor residue pairs previously identified through Cysteine cross-linking studies on our IK_S_ models. Each residue is colored according to the chemical nature of its sidechain. Apolar residues (aliphatic, aromatic) are written in black, basic residues in blue, acidic residues in red, and polar residues in green.

| Residue Pairs |  |  |  | Assigned state | Reference | Number of subunits in 4PIP<sub>2</sub> systems (over 4) |  |  | Number of subunits in 8PIP<sub>2</sub> systems (over 4) |  |  |
| --- | --- | --- | --- | --- | --- | --- | --- | --- | --- | --- | --- |
| KCNE1 region |  | Kv7.1 segment |  |  |  | AO | IC | RC | AO | IC | RC |
| Gly40 | N<sub>TERM</sub> | Thr144 | S1 | AO | (Wang et al., 2011) | 4 | 0 | 1 | 4 | 0 | 3 |
| Gly40 | N<sub>TERM</sub> | Ile145 | S1 |  | (Xu et al., 2008) | 4 | 2 | 3 | 4 | 0 | 2 |
| Gly40 | N<sub>TERM</sub> | Gln147 | S1 |  | (Wang et al., 2011) | 4 | 0 | 1 | 4 | 0 | 2 |
| Lys41 | N<sub>TERM</sub> | Gln147 | S1 |  | (Wang et al., 2011) | 1 | 1 | 3 | 1 | 0 | 3 |
| Leu42 | N<sub>TERM</sub> | Val324 | S6 |  | (Chung et al., 2009) | 4 | 2 | 1 | 4 | 3 | 1 |
| Lys41 | N<sub>TERM</sub> | Thr144 | S1 | RC | (Chung et al., 2009) | 2 | 2 | 4 | 3 | 1 | 3 |
| Lys41 | N<sub>TERM</sub> | Ile145 | S1 |  | (Xu et al., 2008) | 4 | 3 | 4 | 4 | 2 | 3 |
| Lys41 | N<sub>TERM</sub> | Val324 | S6 |  | (Chung et al., 2009) | 0 | 2 | 2 | 0 | 2 | 3 |
| Leu42 | N<sub>TERM</sub> | Ile145 | S1 |  | (Chung et al., 2009) | 4 | 3 | 4 | 4 | 2 | 3 |
| Leu42 | N<sub>TERM</sub> | Gln147 | S1 |  | (Wang et al., 2011) | 0 | 0 | 3 | 0 | 0 | 3 |
| Glu43 | N<sub>TERM</sub> | Gln147 | S1 |  | (Wang et al., 2011) | 2 | 0 | 0 | 1 | 0 | 1 |
| Glu43 | N<sub>TERM</sub> | Trp323 | S6 | Not specified | (Chan et al., 2012) | 4 | 4 | 4 | 4 | 4 | 4 |
| Leu51 | TMD | Cys331 | S6 |  | (Tapper and George, 2001) | 3 | 4 | 4 | 4 | 4 | 4 |
| Glu43 | N<sub>TERM</sub> | Val141 | S1 |  | (Chan et al., 2012) | 4 | 3 | 3 | 4 | 2 | 4 |
| Ala44 | N<sub>TERM</sub> | Val141 | S1 |  | (Chan et al., 2012) | 4 | 3 | 3 | 4 | 3 | 2 |
| Phe54 | TMD | Cys331 | S6 |  | (Tapper and George, 2001) | 4 | 3 | 4 | 4 | 4 | 2 |
| Leu51 | TMD | Ile328 | S6 |  | (Tapper and George, 2001) | 0 | 3 | 4 | 0 | 4 | 4 |
| Ser74 | C<sub>TERM</sub> | His363 | S6 |  | (Lvov et al., 2010) | 0 | 3 | 2 | 0 | 3 | 2 |
| His73 | C<sub>TERM</sub> | His363 | S6 |  | (Lvov et al., 2010) | 0 | 1 | 2 | 0 | 1 | 0 |
| Asp76 | C<sub>TERM</sub> | His363 | S6 |  | (Lvov et al., 2010) | 0 | 1 | 4 | 0 | 1 | 2 |

**Table S4:** State-dependent protein-lipid interactions between K_V_7.1/KCNE1 basic residues and PIP_2_. Identification of the basic residues from K_V_7.1 and KCNE1 subunits which have been previously shown to interact with PIP_2_ through neutralizing point mutations. In the first column, residue names are colored according to its sidechain chemical nature. Apolar residues are colored in black, basic residues in blue, and polar residues in green.

| Mutation | Segment | Reference | Number of subunits in IK<sub>s</sub> models (over 4) |  |  |  |  |  |  |  |  |
| --- | --- | --- | --- | --- | --- | --- | --- | --- | --- | --- | --- |
|  |  |  | PIP2 intra (4PIP<sub>2</sub> systems) |  |  | PIP2 intra (8PIP<sub>2</sub> systems) |  |  | PIP2 inter (8PIP<sub>2</sub> systems) |  |  |
|  |  |  | AO | IC | RC | AO | IC | RC | AO | IC | RC |
| Arg190Cys | S2-S3<sub>LOOP</sub> | (Eckey et al., 2014; Zaydman et al., 2013) | 4 | 4 | 3 | 4 | 4 | 4 | 0 | 0 | 0 |
| Arg195Gln |  |  | 4 | 3 | 3 | 4 | 4 | 2 | 0 | 0 | 0 |
| Arg192Cys |  |  | 2 | 3 | 0 | 3 | 0 | 0 | 0 | 0 | 0 |
| Lys183Cys |  |  | 3 | 0 | 2 | 2 | 0 | 2 | 0 | 0 | 0 |
| Arg243His | S4 | (Park et al., 2005) | 0 | 3 | 4 | 0 | 2 | 4 | 0 | 0 | 0 |
| Arg249Cys | S4-S5<sub>LINKER</sub> | (Eckey et al., 2014; Zaydman et al., 2013) | 3 | 4 | 4 | 2 | 4 | 4 | 1 | 0 | 0 |
| Lys354Cys | S6 |  | 3 | 3 | 1 | 2 | 2 | 0 | 0 | 0 | 0 |
| Lys358Cys |  | (Thomas et al., 2011; Zaydman et al., 2013) | 1 | 2 | 1 | 1 | 0 | 0 | 3 | 2 | 1 |
| Lys362Cys |  |  | 0 | 0 | 0 | 0 | 0 | 0 | 4 | 4 | 3 |
| Arg366Ala |  |  | 1 | 0 | 1 | 0 | 0 | 0 | 4 | 4 | 3 |
| Arg67Gln | KCNE1<sub>M</sub><sup>CTER</sup> | (Li et al., 2011) | 1 | 0 | 0 | 1 | 0 | 0 | 0 | 4 | 4 |
| Lys69Cys |  |  | 0 | 4 | 2 | 0 | 4 | 2 | 4 | 0 | 0 |
| Lys70Cys |  |  | 0 | 0 | 0 | 0 | 1 | 0 | 0 | 1 | 3 |

**Figure S1:**
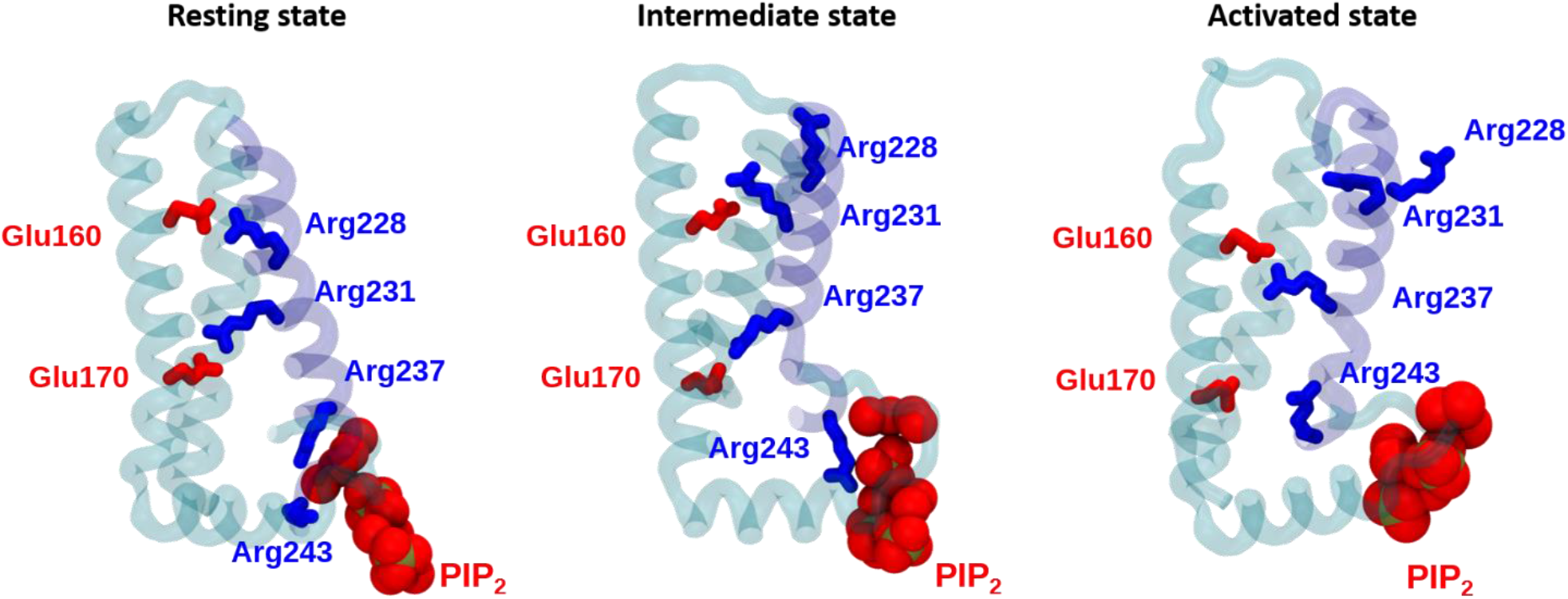
State-dependent ionic interactions in the VSD of K_V_7.1 subunits. Mapping of S4 gating charges (in blue sticks) along with their VSD countercharges (in red sticks) and PIP_2_ intra phosphate groups (in red spheres) in stable state models of IK_S_ and K_V_7.1 channels. VSD segments are represented in transparent cyan ribbons, except S4 (in transparent blue ribbons) and S3 segment, hidden for clarity.

**Figure S2:**
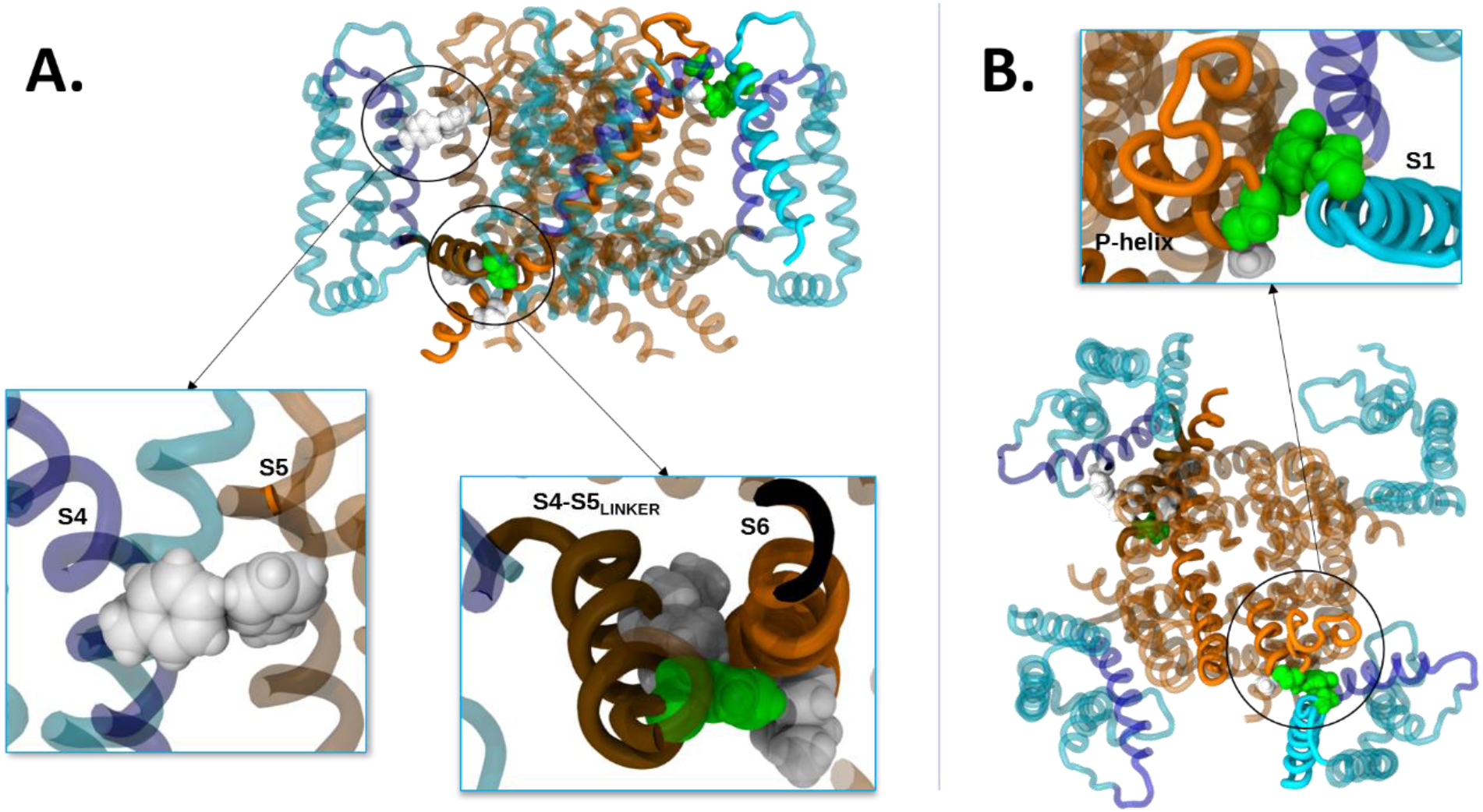
State-dependent K_V_7.1 protein-protein interactions in IK_S_ models. Structural mapping of state-dependent protein-protein interactions in our IK_S_ models in **A.** Side view and **B.** Top view. KCNE1 segments are hidden for clarity. K_V_7.1 segments are represented in transparent ribbons, except those which carry the residue pairs of interest, which are in solid colors. Neighbor residue pairs are circled in black, and link to a zoomed view. For transmembrane segments, VSD ones are colored in cyan, S4 in blue, S4-S5_LINKER_ in brown and PD segments in orange. Residues of interest are colored according to the chemical nature of their sidechains: apolar residues are colored in white, and polar residues are in green

**Figure S3:**
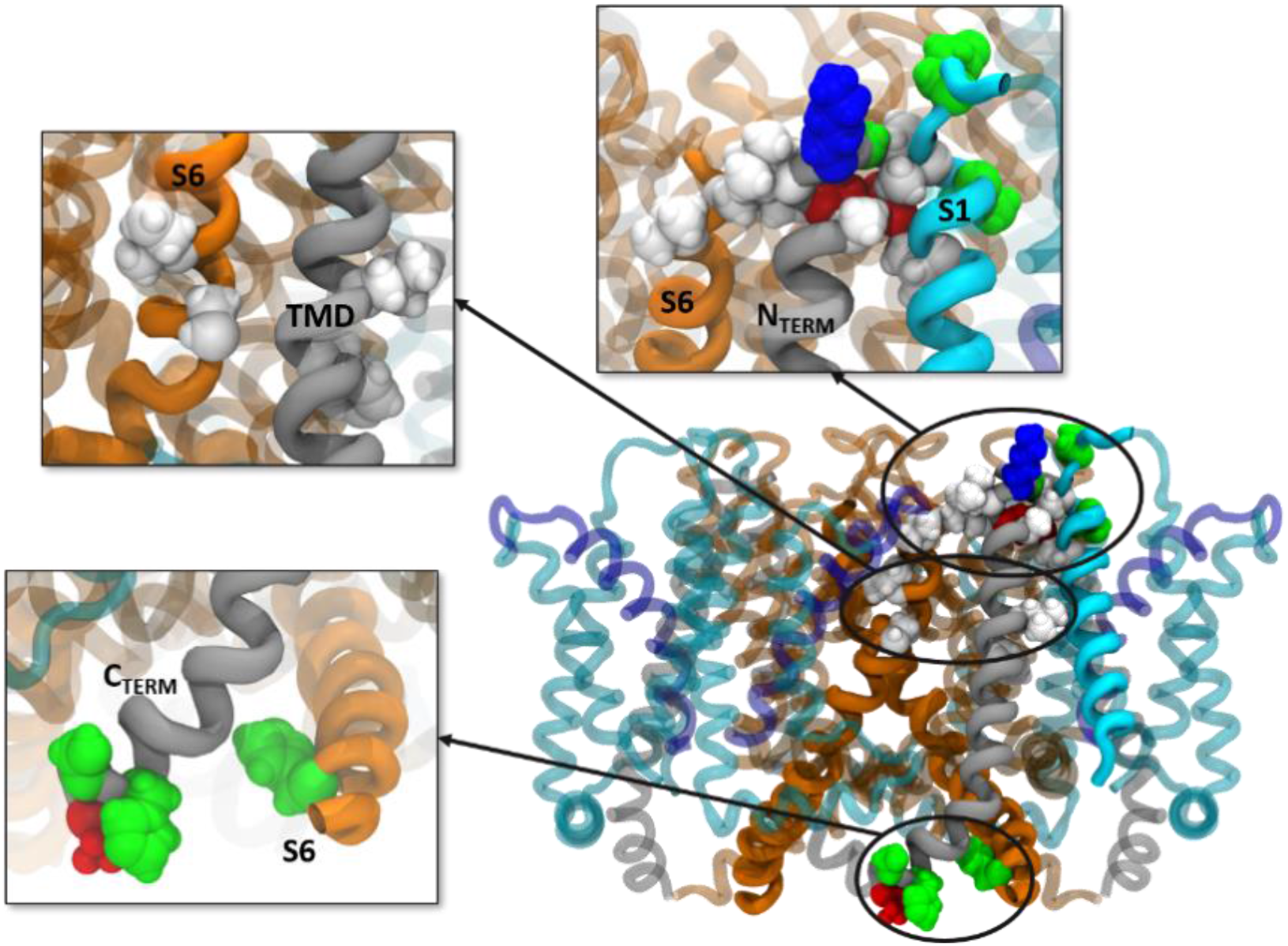
State-dependent KCNE1-K_V_7.1 interactions in IK_S_ models. Structural mapping of state-dependent protein-protein interactions in IK_S_ models. KCNE1 segments are represented in grey ribbons. K_V_7.1 segments are represented in transparent ribbons, except those containing the residue pairs of interest, which are in solid colors. Neighbor residue pairs are circled in black, and for which a zoomed view is provided. For transmembrane segments, the color code used is the same as those of Figure S2. Residues of interest are colored according to the chemical nature of their sidechains: apolar residues are colored in white, acidic residues in red, basic residues in blue, and polar residues in green.

**Figure S4:**
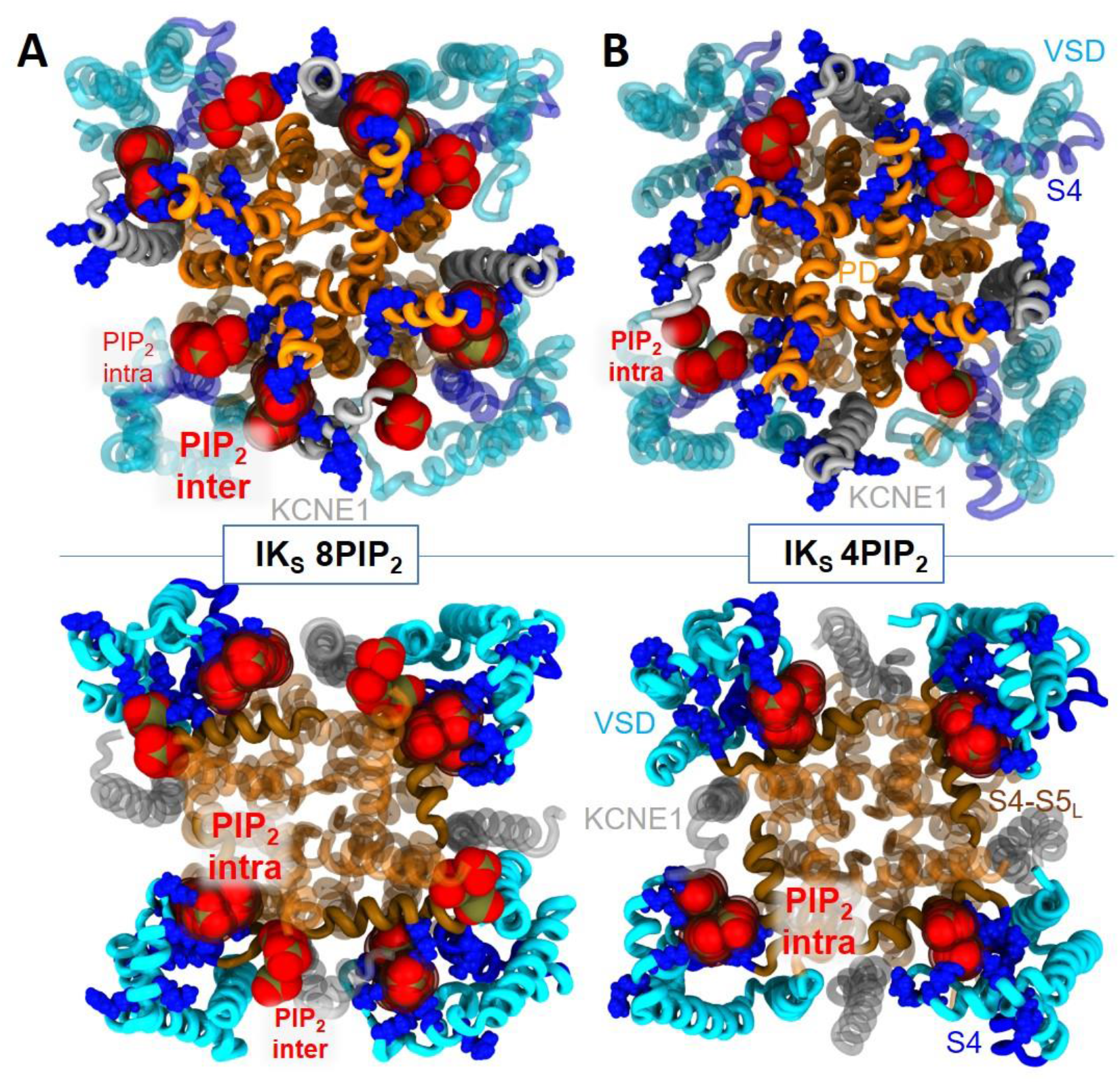
Structural mapping of the salt-bridges between IK_S_ complex residues and PIP_2_. Cytoplasmic view of the transmembrane subunits (in ribbons) of the AO models of IK_S_ complex. VSD segments are colored in cyan, S4 segments in blue, PD segments in orange and KCNE1 subunits in gray. In each panel, the basic residues of IK_S_ complex are depicted in blue spheres, while the charge moieties of PIP_2_ inter and PIP_2_ intra are depicted in bright red and dark red spheres, respectively. **A.** In 8PIP_2_ systems, the upper panel shows the cytoplasmic regions of KCNE1 and S6 (in solid colors) which specifically interact with PIP_2_ inter. The lower panel shows the cytoplasmic regions of the VSD including S2-S3_LOOP_, S4 and S4-S5_LINKER_ (in solid colors) that interact with PIP_2_ intra. **B.** In 4PIP_2_ systems, the upper panel depicts the relative position of the basic residues of KCNE1 (in solid colors) with respect to PIP_2_ intra that interacts with the cytoplasmic region of S6, as PIP_2_ inter lipids are absent. The lower panel shows the interactions between the basic residues from S2-S3_LOOP_, S4 and S4-S5_LINKER_ (in solid colors) that interact with PIP_2_ intra (in circled spheres).

